# Network-based genetic monitoring of landscape fragmentation

**DOI:** 10.1101/2025.06.18.660081

**Authors:** Ohad Peled, Jaehee Kim, Gili Greenbaum

## Abstract

Habitat fragmentation is one of the most immediate and substantial threats to biodiversity, generating isolated populations with reduced genetic diversity. Genetic monitoring has become essential for detecting fragmentation and tracking its progress. However, the coherent interpretation of genetic monitoring data and understanding the genetic consequences of fragmentation require frameworks that accurately represent real-world complexity. Existing theoretical frameworks typically rely on simplified spatial structures and do not adequately capture the heterogeneous migration patterns of natural populations. Here, we integrate network theory and mathematical population genetics to develop a framework for studying the genetic consequences of fragmentation processes, explicitly accounting for heterogeneous connectivity and temporal dynamics. We apply this framework to examine how different fragmentation processes affect genetic measures commonly used in genetic monitoring. We find that different fragmentation scenarios produce substantially distinct trajectories in key genetic measures, sometimes exhibiting rapid transitional dynamics, suggesting that the interpretation of genetic monitoring data must be tailored to ecological contexts. Furthermore, fragmentation can cause deviations from classical theoretical expectations, such as the expected correlation between genetic and geographic distance (isolation-by-distance) or between genetic diversity and connectivity. Finally, we propose and demonstrate detectable early warning signals in genetic monitoring data that precede rapid transitional phases. Our framework thus provides a practical interpretation of genetic monitoring data, bridging the gap between idealized theoretical models and real-world connectivity dynamics.

## Introduction

Rapid human-induced environmental changes affect ecological and evolutionary processes, driving biodiversity loss [1]. One of the main factors driving these changes is landscape fragmentation, the partitioning of landscapes into small and weakly connected habitat patches [2]. Fragmentation reduces connectivity among populations, constraining gene flow and dispersal of individuals [3], which can negatively impact the health and viability of populations [4–6]. Landscape fragmentation is expected to erode within-population genetic diversity and increase between-population genetic differentiation due to reduced gene flow and increased genetic drift [7, 8]. Decreased genetic diversity can, in turn, reduce population viability in the short term by increasing risks of inbreeding depression [7, 9], while also limiting long-term evolutionary potential and adaptive capacity in response to future environmental changes [10, 11]. Consequently, systematically and coherently tracking fragmentation dynamics and their population-genetic consequences through genetic monitoring remains a major goal in conservation biology.

Genetic monitoring of population genetic metrics over time is a cost-effective and direct approach for tracking both the genetic impacts and the underlying ecological processes of fragmentation. The alternative, tracking individual movement among habitat patches, is usually resource-intensive and offers only an indirect proxy for the genetic and evolutionary consequences of fragmentation. Consequently, genetic monitoring of wild populations is widely used to assess population health and viability, landscape connectivity, and species responses to environmental disturbances [12–14]. However, a major challenge in applying genetic monitoring to track fragmentation lies in the interpretation of genetic measures in the context of the ecological process of migration.

Early theoretical work in population genetics established foundational frameworks for linking genetic diversity and differentiation to migration under simplified assumptions about gene flow patterns and spatial configurations [15–17]. For example, the island model assumes equal and constant migration rates without explicit spatial arrangement [15], whereas the stepping-stone model incorporates homogeneous and symmetric migration between adjacent demes arranged on a regular lattice with an additional long-range migration component [17]. These models provided fundamental insights into how spatial connectivity shapes population genetic structure and introduced key concepts such as isolation-by-distance, where genetic differentiation increases with geographic distance [18], and the connectivity-diversity relationship, in which populations that are more connected are expected to exhibit higher genetic diversity [19]. However, their simplified assumptions often limit their practical applicability for genetic monitoring and evaluating fragmentation impacts [20]. For example, one critical limitation of most existing modeling frameworks is their inability to capture the temporal dynamics of fragmentation, where landscape degradation and connectivity loss occur as heterogeneous, sequential processes shaped by the specific spatial and temporal characteristics of anthropogenic or climatic drivers [21]. The lack of a modeling framework that integrates realistic spatiotemporal patterns of connectivity and fragmentation thus restricts the practical application of population genetic theory in conservation efforts and limits its utility for informing management decisions.

A promising approach for incorporating realistic gene flow patterns into population genetic theory is to represent connectivity between populations as a network—a mathematical construct comprising nodes (habitat patches) connected by edges (connectivity) [22]. Population networks can accommodate complex connectivity patterns beyond the scope of classical population genetics models. Several methods have been developed to infer such networks from genetic data by quantifying genetic differentiation between population pairs [23–25], with applications across a wide range of taxa [23, 26–32]. These network-based approaches provide a rigorous framework for modeling realistic fragmentation dynamics, enabling more coherent interpretations of genetic monitoring.

In this work, to bridge the gap between theory and practice, we develop a framework based on population networks, integrating advances in population-genetic theory and network science to investigate the spatiotemporal genetic consequences of landscape fragmentation. This framework explicitly incorporates real-world complexities within a conceptually simple and tractable model. We apply this framework to examine how different fragmentation scenarios affect genetic measures and to assess how network structure impacts population resilience under connectivity loss. While fragmentation is a multifaceted process involving multiple concurrent stressors (e.g., habitat loss, reduced patch size, edge effects), our focus here is on connectivity loss (also termed fragmentation *per se*; [5, 33]). Our approach enables improved interpretation of genetic monitoring data and facilitates identification and measurement of fragmentation progression. Additionally, our modeling framework can assist in predicting the genetic impacts of connectivity loss and evaluating the genetic health of fragmented populations.

## Results

To model the genetic consequences of fragmentation, we consider a metapopulation in which some populations are connected by migration. For tractability, we assume equal and symmetric migration rates among all connected populations. Any such connectivity pattern can be represented as a population network (Fig. 1a). To relate migration patterns to genetic measures, we employ the approach developed by Alcala *et al.* [34], which consists of two transformations: (i) from migration matrices to pairwise coalescent-time matrices [35], and (ii) from coalescent-time matrices to pairwise genetic differentiation measured by *F_ST_* [36] (see *Methods* and Supplementary Information Text). This procedure provides, for a given migration matrix, expected pairwise *F_ST_* between all population pairs, as well as genetic diversity measured by expected heterozygosity (*H_e_*) for each population (Fig. 1a). For simplicity, we further assume uniform population sizes and mutation rates across all populations, allowing us to use an ‘unscaled’ heterozygosity measure (see *Methods*); therefore, our *H_e_* values should be interpreted only relatively, and values exceeding one are possible.

**Figure 1:**
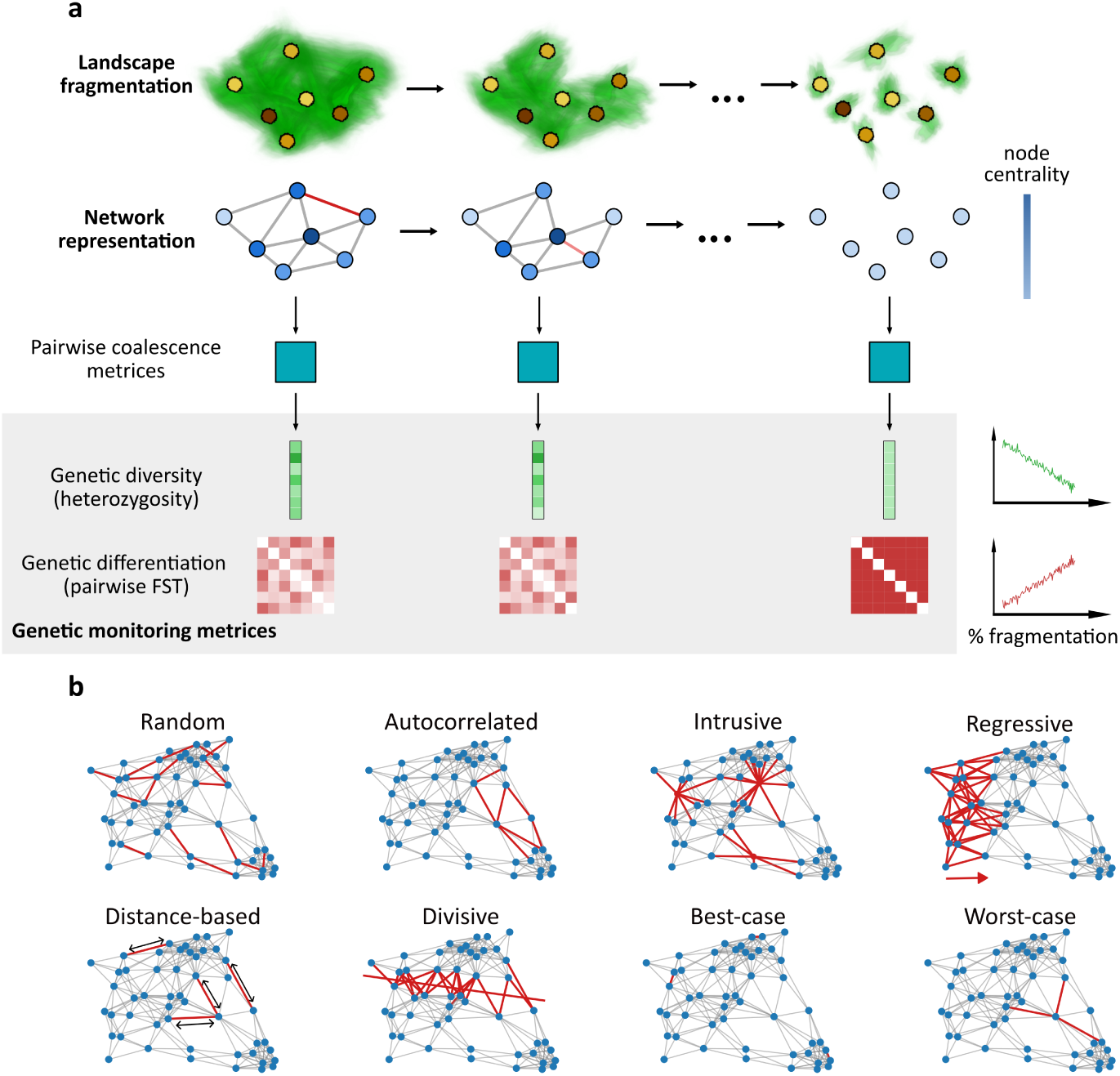
Schematic representation of the network-based framework for modeling population genetic effects of fragmentation. (a) Computation of genetic measures along fragmentation. In the top row, populations (yellow/brown patches) are embedded in a landscape (green) undergoing fragmentation. Below, the metapopulation is represented as a network with nodes (blue) denoting populations and edges representing migration between populations. Fragmentation is simulated by iteratively removing edges (red). A coalescence matrix is derived from each network, which enables the calculation of genetic diversity and differentiation at each fragmentation step (grey box). These metrics allow monitoring of population genetic changes over time (right side of the grey box). Color intensity of nodes represents network properties associated with genetic measures. (b) Modeling fragmentation processes. Illustrated are eight fragmentation scenarios applied to a single realization of a random geometric graph (RGG). Edges removed under each scenario are shown in red. Further details of each scenario are provided in the text.

To simulate an ecologically plausible metapopulation, which is usually embedded in a geographic landscape, we use a random geometric graph (RGG) model [37] as the initial network. In this model, populations are more likely to be connected if they are geographically close to each other. We model a fragmentation process by iteratively removing edges according to one of several predefined fragmentation scenarios (Fig. 1b). After each edge removal, we recompute genetic measures, tracking their changes until all edges have been removed and the network has become fully fragmented into isolated populations. This modeling framework is highly flexible and enables the study of diverse connectivity patterns and fragmentation scenarios while providing rigorous analytical expectations for key genetic measures commonly used in genetic monitoring.

We consider eight fragmentation scenarios (Fig. 1b): (i) random fragmentation, representing global environmental changes (e.g., climate change); (ii) autocorrelated fragmentation, representing spatially correlated landscape disturbances (e.g., agricultural expansion); (iii) intrusive fragmentation, representing the emergence of isolated habitats within the landscape; (iv) regressive fragmentation, representing the expansion of a disturbance into a natural landscape (e.g., urban expansion);(v) distance-based fragmentation, representing reduced dispersal ability through a non-habitable matrix (e.g., disturbances hindering dispersal through the matrix, reducing dispersal distances); (vi) divisive fragmentation, representing linear destruction of connectivity (e.g., road or railway construction); (vii) best-case fragmentation, an idealized scenario that sequentially removes the least important edges, thus maximizing connectivity at each step; and (viii) worst-case fragmentation, similar to the best-case scenario, except the most important edge is removed at each step. The last two scenarios are theoretical constructs intended to establish upper and lower bounds for genetic measures rather than to depict realistic fragmentation processes. Detailed descriptions of each fragmentation scenario are provided in *Methods*.

### Genetic monitoring measures strongly depend on the fragmentation scenario

Across all fragmentation scenarios, we observe an increase in genetic differentiation and a decrease in genetic diversity as fragmentation progresses (Fig. 2). However, the rate and pattern of these changes vary substantially among scenarios. The slowest erosion of genetic diversity and the most gradual increase in genetic differentiation were observed under the best-case scenario (pink curve in Fig. 2), as expected. In contrast, the worst-case scenario exhibited the most rapid erosion of genetic diversity and the steepest increase in differentiation (grey curve in Fig. 2). Thus, these two theoretical extremes provide upper and lower bounds for the retention of genetic health in the metapopulation, against which other fragmentation scenarios can be compared.

**Figure 2:**
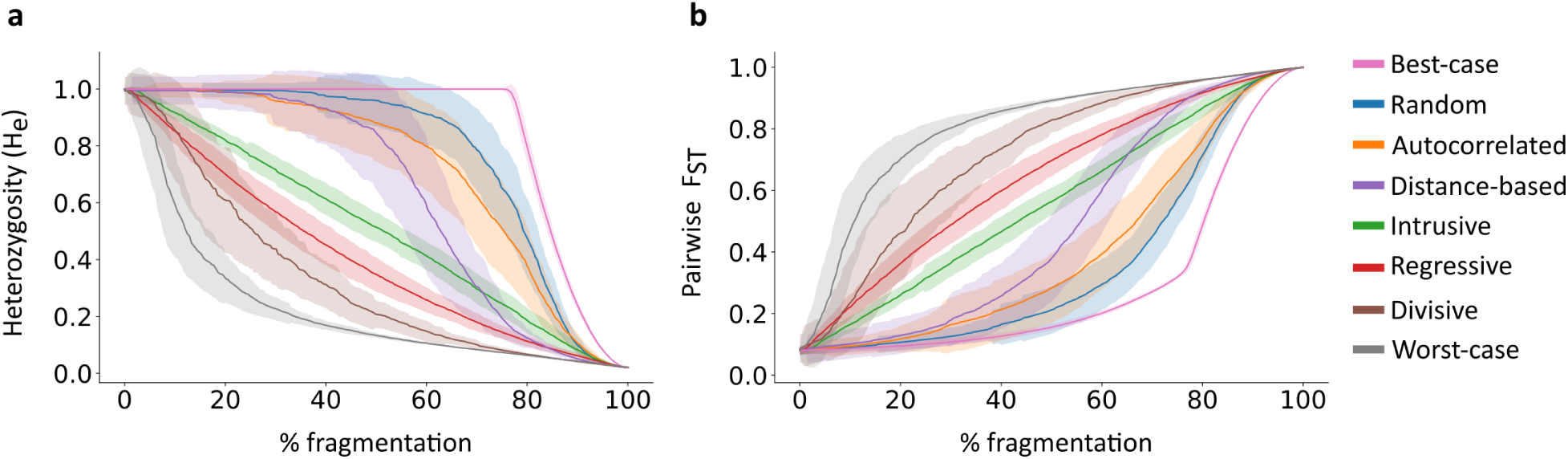
Changes in genetic measures along fragmentation under eight fragmentation scenarios. (a) Mean genetic diversity (*H_e_*) across all populations along fragmentation. (b) Mean genetic differentiation (pairwise *F_ST_*) among all population pairs. Lines denote means across 100 simulation replicates, with shaded regions indicating standard deviations. Fragmentation is measured as the fraction of edges removed from the initial network.

In the random and autocorrelated scenarios, the loss of diversity and increase in differentiation are almost undetectable in the early stages of fragmentation but then become substantial at ∼ 60% fragmentation. This pattern is reflected in concave curves for genetic diversity and convex curves for differentiation (blue and orange curves in Fig. 2). The distance-based scenario (purple curve in Fig. 2) shows a similar trend, but the loss of genetic diversity begins earlier in the fragmentation process and progresses faster than in the random and autocorrelated scenarios. In contrast, in the regressive and divisive scenarios, the curvature patterns are reversed: the genetic diversity curve is convex, with rapid and substantial decreases in genetic diversity early in the fragmentation process, and the genetic differentiation curve is concave, indicating earlier deterioration of metapopulation genetic health compared to the other scenarios. For example, in the divisive scenario, a *>* 50% change in genetic measures occurs already by 25% of the fragmentation process (brown curve in Fig. 2). In the intrusive scenario, both genetic measures change approximately linearly as fragmentation progresses (green curve in Fig. 2).

To understand the robustness of these patterns, we also examined how *F_ST_* and *H_e_* measures change along fragmentation under different migration rates and initial network topologies (Figs. S1 and S2). Overall, the patterns remain similar across different migration rates, except at low migration rates, where the absolute values of *F_ST_* are higher in the early stages of fragmentation (Fig. S1b). Similarly, the results remained consistent when the initial network topologies were generated using either the Erdős-Ŕenyi model or a small-world network model instead of the RGG model (Fig. S2). However, the differences among fragmentation scenarios were less pronounced in these analyses, highlighting the importance of considering spatially explicit network models, such as the RGG model.

Overall, our results demonstrate that, for a given level of connectivity loss, the risk of inbreeding depression and the reductions in both evolutionary potential and between-population differentiation strongly depend on the type of fragmentation process experienced by the metapopulation. Therefore, the interpretation of genetic monitoring data must account for the context and drivers of fragmentation. For example, a 10% decrease in *H_e_* might reflect gradual connectivity decline under intrusive fragmentation, whereas the same decrease under random fragmentation could indicate dramatic habitat deterioration.

### Relationship between heterozygosity and network components

When considering the distributions of the genetic measures rather than just their means, we observe that *H_e_* distributions remain largely unimodal throughout the fragmentation process, with a shift towards *H_e_* = 0 occurring as isolated nodes accumulate (Figs. 3a–c and S4a–e). Similarly, the *F_ST_* distributions exhibit increasing bimodality, with density accumulating at *F_ST_* = 1 as more nodes are separated into different components (Figs. 3d–f and S4f–j). Changes in the shape of these distributions along fragmentation are also reflected in the variance of genetic diversity across populations (Fig. 3g): the level of fragmentation that maximizes variance, as well as the maximum variance value, differs among fragmentation scenarios. The increase in *H_e_* variance can make the detection of fragmentation—and genetic health in general—more challenging at intermediate fragmentation levels because more populations will need to be sampled to correctly characterize the genetic diversity state of the metapopulation.

**Figure 3:**
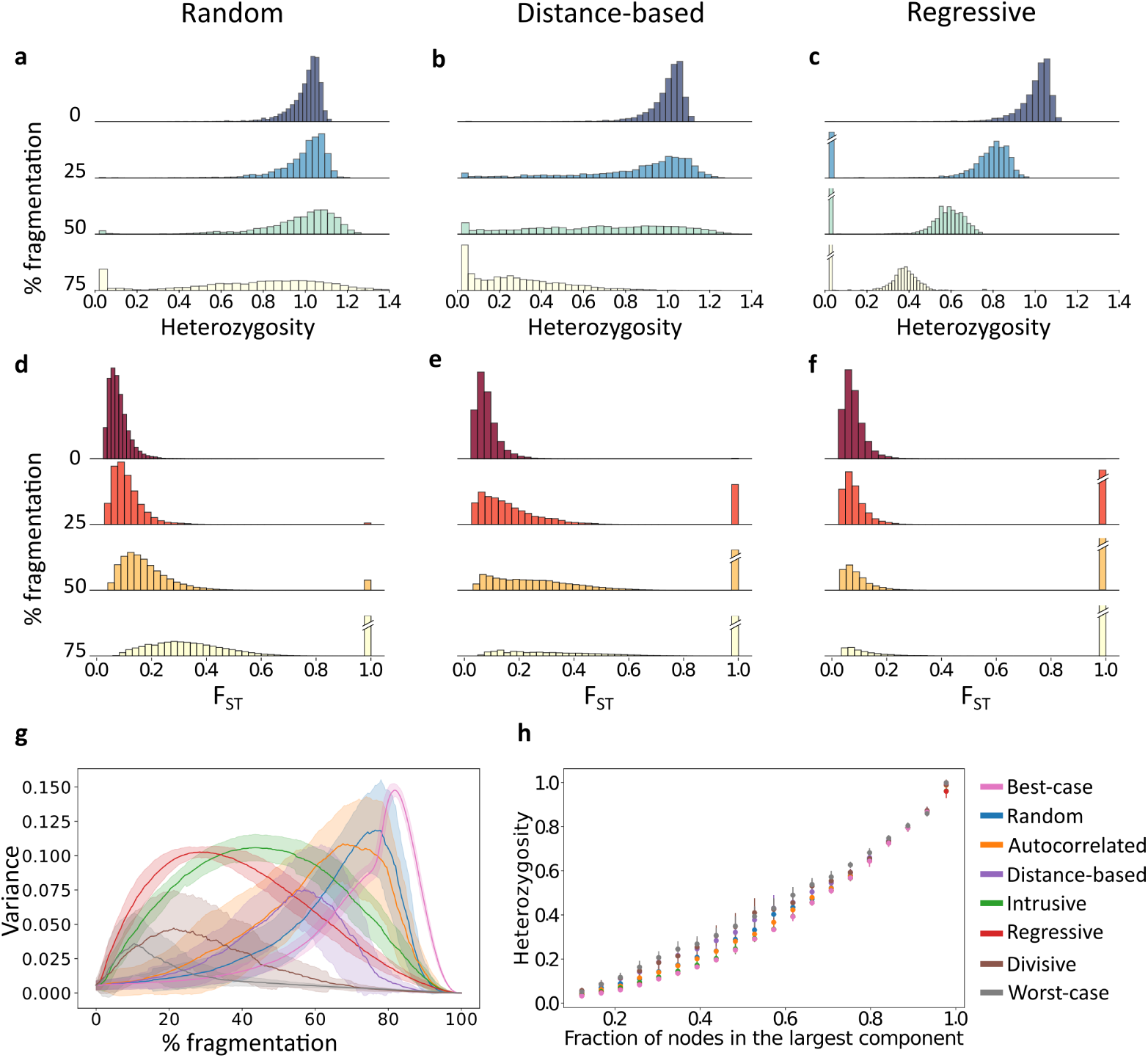
Changes in the distributions of genetic measures along fragmentation. Panels (a–f) show density distributions for three fragmentation scenarios: random, distance-based, and regressive (additional fragmentation scenarios are shown in Fig. S4). Four snapshots from the process are shown: 0%, 25%, 50%, and 75% fragmentation. Diagonal lines on bars indicate truncated values (for *H_e_* = 0 or *F_ST_*= 1). All distributions are pooled from 100 simulation replicates. (a–c) Distribution of expected heterozygosity (*H_e_*) of populations. (d–f) Distribution of pairwise *F_ST_* across all population pairs. (g) Change in the variance of *H_e_* across all populations in the network. (h) Relationship between the fraction of nodes in the largest component and mean *H_e_* across all populations in each network. For each scenario, dots denote the means across 100 simulation replicates, and lines denote the standard deviations.

As fragmentation progresses, network structure changes and populations begin to disconnect from the main component (Fig. S3). For example, the rapid deterioration in genetic health under the divisive scenario (brown in Fig. 2) can be attributed to the early emergence of medium and small network components, which reduce genetic diversity and increase between-component differentiation (Fig. S3f). To better understand the effect of component structure on genetic diversity, we tracked the size of the largest component throughout the fragmentation process (Fig. 3h). We observe a strong correlation between the size of the largest component and the mean *H_e_* across populations in the network (*r* = 0.97–0.98 across scenarios, *p*-value *<* 0.001). This correlation is relatively consistent across different fragmentation scenarios, indicating that the size of the largest component is an important determinant of genetic diversity.

This result can be interpreted in relation to the theoretical relationship between effective population size and heterozygosity, 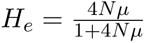 [38]. Because we consider a small effective population size relative to the mutation rate (i.e., *θ* = 4*Nµ* ≪ 1), we expect an approximately linear relationship of *H_e_* ≈ 4*Nµ*. The result in Fig. 3h is similar to what one would expect if we treated each component as a well-mixed population. However, the relationship between *H_e_* and component size is sublinear, reflecting the fact that components are not well-mixed and should therefore be represented with effective sizes smaller than their actual sizes.

### Using network metrics in genetic monitoring

To better understand how tracking network characteristics can inform genetic monitoring, we evaluated the association between genetic measures and commonly used network metrics. We first examined the relationship between a population’s genetic diversity and its centrality. There are different ways to measure network centrality [39], each of which can be interpreted differently with respect to population genetic processes [22]. Here, we evaluated two common metrics: degree centrality (i.e., the number of edges incident to a node), which measures local centrality, and betweenness centrality (i.e., the frequency with which a node lies on shortest paths between other nodes), which measures global centrality. Under classical population genetics theory, populations with higher connectivity should exhibit greater genetic diversity due to increased gene flow, leading to higher *H_e_* at migration-drift equilibrium [19]. Consistent with this expectation, analysis of the initial (pre-fragmentation) networks showed a strong positive correlation between degree centrality and *H_e_*(*r* = 0.71–0.95, Fig. 4a). However, because all populations had a relatively high *H_e_*, this relationship was nonlinear, exhibiting a saturating effect: while *H_e_* increased with degree at low connectivity, it plateaued for highly connected nodes (Fig. S5a). Hence, local connectivity increases genetic diversity only up to a threshold, beyond which additional migration corridors do not significantly contribute to maintaining genetic diversity. In contrast, the association between *H_e_* and betweenness centrality was weaker for nodes with low betweenness (Fig. S5b).

**Figure 4:**
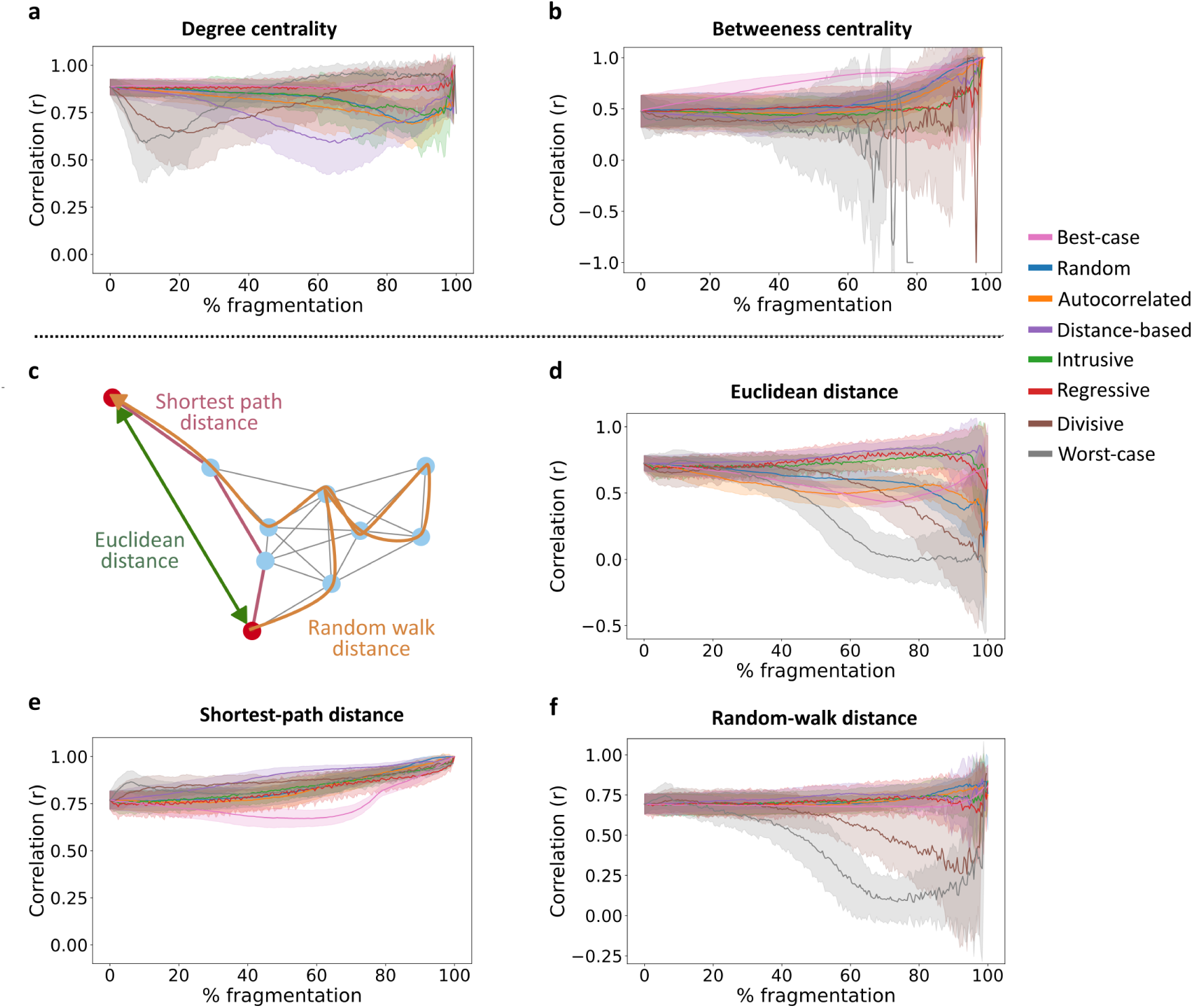
Correlation between population genetic measures and network metrics. The Pearson correlation coefficient *r* was computed between genetic diversity (*H_e_*) and network centrality (panels a–b), or between genetic differentiation (*F_ST_*) and distance metrics (panels c–f), for eight fragmentation scenarios. (a) Correlation between a population’s *H_e_* and its degree centrality (number of connected edges). (b) Correlation between a population’s *H_e_* and its betweenness centrality (global centrality metric). (c) Schematic illustration of three different distance metrics for a pair of populations (red nodes). (d) Correlation between the *F_ST_* of a pair of populations and their Euclidean distance in the two-dimensional space in which the RGG network is embedded. (e) Correlation between the *F_ST_* of a pair of populations and their shortest-path network distance. (f) Correlation between the *F_ST_* of a pair of populations and their random-walk network distance.

Throughout fragmentation, the correlation between *H_e_* and degree centrality remains consistently high for some scenarios but declines rapidly early in the fragmentation process under the worst-case, divisive, and distance-based scenarios (Fig. 4a). This decline may result from network partitioning into components of varying size in these fragmentation scenarios, where component size has a stronger effect on *H_e_* than does local connectivity. For example, a densely connected population in a small component with few populations may have lower *H_e_* than a sparsely connected population in a larger component with many populations. Thus, component size, rather than degree centrality, is a primary determinant of genetic diversity at these intermediate fragmentation stages. Interestingly, in these scenarios, the correlation later rebounds, converging to levels similar to those of the other fragmentation scenarios. This suggests that once components reach comparatively small sizes, within-component degree centrality once again becomes a strong determinant of *H_e_*.

The association between genetic diversity and betweenness centrality was generally weaker than that for degree centrality, with less variation among fragmentation scenarios (Fig. 4b). This suggests that populations do not necessarily need to occupy a key gene flow hub to maintain high genetic diversity, as has been observed in some systems [40]. One implication of this is that peripheral populations in large, well-connected networks can maintain genetic diversity comparable to that of central populations in smaller, less connected components.

Next, we examined the relationship between pairwise *F_ST_* and three network distance metrics relevant for genetic monitoring (Fig. 4c): (i) Euclidean distance in the two-dimensional space of the embedded RGG network, (ii) shortest-path distance (the minimum number of edges required to connect a pair of nodes), and (iii) random-walk distance (the expected number of edges traversed in a random walk between two nodes). Euclidean distance represents the geographic distance, whereas estimating the network distances requires knowledge of migration or movement patterns in the metapopulation [22]. Prior to fragmentation, we observed strong correlations between *F_ST_* and all three distance metrics (*r* = 0.7–0.75, Fig. 4d–f), indicating that geographically distant populations are more genetically differentiated irrespective of how distance is measured. This finding aligns with the isolation-by-distance expectation derived from stepping-stone [17] and continuous-space [41] models, suggesting that such idealized models are a good approximation for sufficiently well-connected networks [22]. However, as fragmentation progresses, these correlations vary substantially both over the course of fragmentation and among scenarios (Fig. 4d–f).

For Euclidean distance, the correlation declines under most fragmentation scenarios, particularly under the worst-case, divisive, best-case, and autocorrelated scenarios (Fig. 4d). Interestingly, these processes differ substantially in their network structure during fragmentation, particularly in the size of the largest connected component (Fig. S3), indicating that additional topological properties are needed to account for the relationship between genetic differentiation and geographic distance. In contrast, for shortest-path distance, the correlation consistently increases across most processes we examined (the theoretical best-case scenario is the exception) and generally shows the strongest association between *F_ST_* and distance (Fig. 4e). This suggests that this network metric is particularly suited for genetic monitoring, either as a non-genetic proxy for genetic differentiation or as a proxy for connectivity from pairwise *F_ST_* data. For the random-walk distance, the relationship remains relatively stable throughout fragmentation for most scenarios, except for a decline in the worst-case and divisive scenarios (Fig. 4f).

Overall, these analyses highlight that the topological properties of population networks can inform the tracking of genetic diversity and differentiation patterns. However, relating genetic measures to network properties such as components, centrality, or distance measures should, in most cases, be done in the context of the fragmentation scenario. Classical population genetic relationships—such as those between gene flow and diversity or distance and differentiation—are useful for well-connected populations but may diverge from classical theory when fragmentation processes shape the topology of metapopulation connectivity.

### Early warning signals in genetic monitoring

The goal of genetic monitoring is to track the genetic health of populations and to infer underlying ecological processes. However, our findings suggest that inferring fragmentation solely from genetic metrics can be challenging because substantial shifts in genetic measures often occur only in the later stages of fragmentation under certain fragmentation scenarios. In such cases, once genetic diversity declines and population differentiation increases, the transition is both rapid and pronounced (Fig. 2). This transition can be considered a tipping-point phase, before which it is difficult to detect ongoing fragmentation by tracking the means of *H_e_* and *F_ST_*. This raises the question: Can genetic monitoring data detect landscape fragmentation early enough—before the population transitions to a highly fragmented and diversity-depleted state? In other words, if we are tracking genetic measures in a metapopulation that is progressively undergoing fragmentation, can we use genetic data to provide an early warning signal prior to the tipping-point phase during which genetic diversity and differentiation dramatically change? To address this question, we evaluated whether early warning signals can be extracted from genetic diversity measures, borrowing methods from complex systems theory [42, 43]. Our analysis provides proof-of-concept for the potential to integrate early warning methodologies into genetic monitoring frameworks.

For this demonstrative analysis, we focused on the genetic diversity under the autocorrelated fragmentation scenario, where edges are removed in a spatially coordinated manner. We first considered a genetic monitoring scheme that tracks the *H_e_* distributions of all populations throughout fragmentation (Fig. 5a). At each time step, we analyzed the distribution of *H_e_* across populations and computed several summary statistics—standard deviation, skewness, and kurtosis—which have been found to be reliable early warning indicators in other disciplines [44, 45]. Another common statistic, lag-1 autocorrelation, was not used because it is intended to measure stability around a single equilibrium [44, 46], which did not hold in our simulations. As the metapopulation approaches the tipping-point phase, the theoretical expectation is that the standard deviation of the *H_e_* distribution will increase, the skewness will shift toward the new state (in this case, asymmetry towards lower *H_e_* values), and the kurtosis will increase due to an increased frequency of extreme values [43, 45]. To evaluate this, we computed these summary statistics throughout fragmentation (green curves in Fig. 5b–d) and examined whether they show substantial changes prior to the tipping-point phase (the sharp drop in the orange curves at ∼ 80–90% fragmentation in Fig. 5b–d).

**Figure 5:**
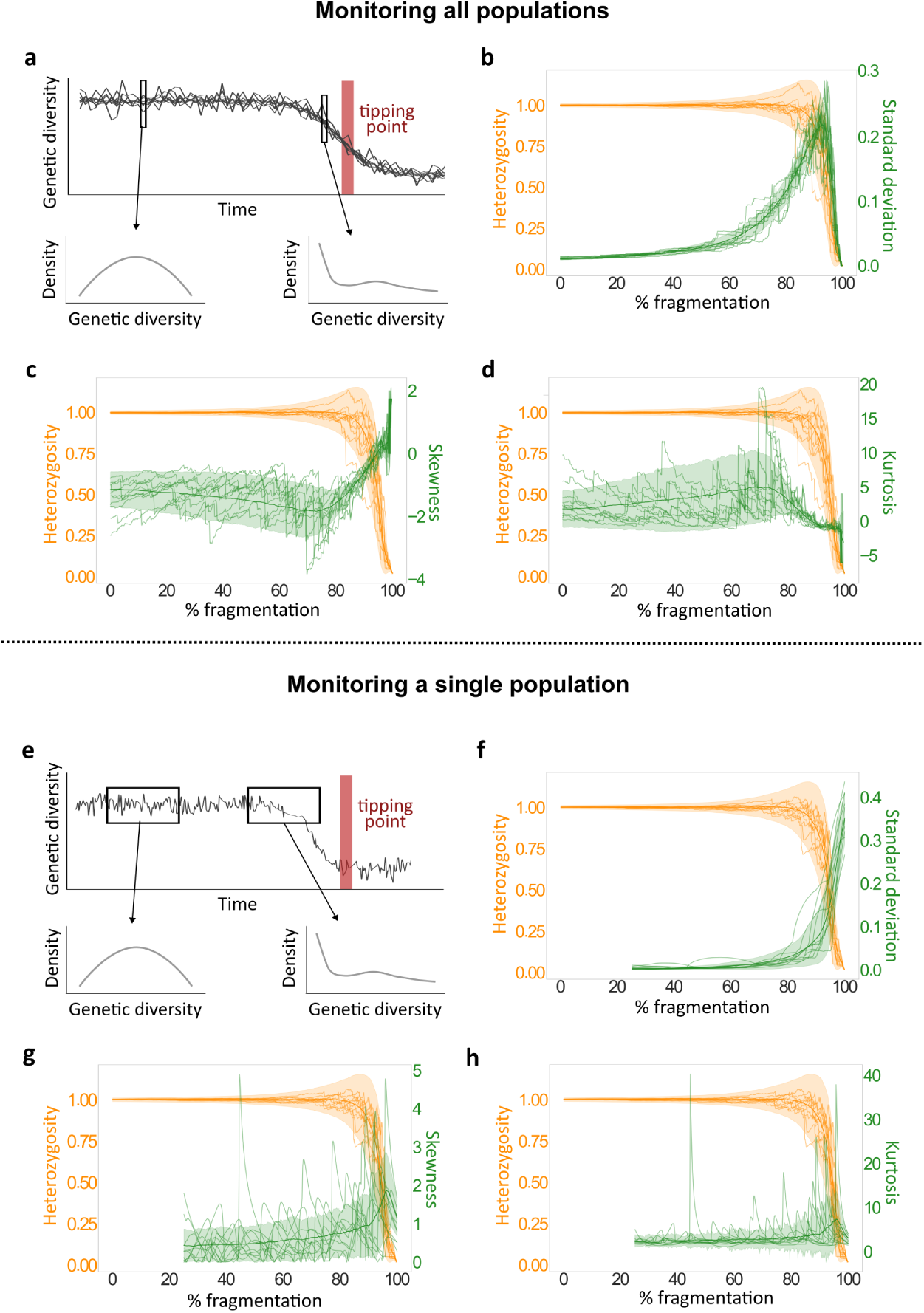
Early warning signals before tipping point in genetic monitoring. The analysis examines fragmentation under the autocorrelated scenario (Fig. 1b). (a) Schematic of the metapopulation-monitoring approach. At each time step, we analyze the *H_e_* distribution of all connected populations in the network (largest component). The tipping-point phase, during which genetic diversity dramatically declines, is denoted in red. Genetic diversity distributions closer to the tipping-point phase may differ, with summary statistics potentially providing early warning. (b–d) Mean metapopulation heterozygosity (orange) and three early warning statistics (green: SD in (b), skewness in (c), and kurtosis in (d)) along fragmentation. Solid lines show the mean across 1000 simulation replicates, shaded areas show the standard deviation, and thin lines show ten individual replicates. (e) Schematic of the single-population monitoring approach. The *H_e_* of a single population is tracked with a sliding window; the *H_e_* distributions in each window are then analyzed. The tipping-point phase is shown in red. As above, the genetic diversity distributions in windows closer to the tipping-point phase may differ, with summary statistics potentially providing an early warning. (f–h) Mean metapopulation heterozygosity (orange) and three early warning statistics computed from 25% of the data per window (green: SD in (f), skewness in (g), and kurtosis in (h)). Solid lines show the mean across 1000 simulation replicates, shaded areas show the standard deviation, and thin lines show ten individual replicates.

Some early warning signals prior to the tipping-point phase were clearly observable in our analyses (Fig. 5b–d). For example, the standard deviation of the distribution of *H_e_* across the metapopulation increases steadily as fragmentation progresses, and substantial changes in this statistic are observable at the early stages of fragmentation even when changes in the mean are not yet detected (Fig. 5b). Thus, by tracking the standard deviation among populations over time, a noticeable change in this summary statistic could be identified and used as an early warning signal before the tipping-point phase. The mean skewness and kurtosis also showed early changes that can serve as early warning signals (Fig. 5c–d). However, the trajectories of skewness and kurtosis fluctuated over time and were noisier than the standard deviation, suggesting that they are less reliable as early warning indicators. This means that, while tracking the mean metapopulation *H_e_* would not indicate that a rapid reduction in genetic health is approaching, monitoring higher moments could potentially provide an early indication of genetic deterioration.

We also considered a more limited monitoring scenario, where *H_e_* is monitored for a single population (Fig. 5e). In this setting, a single population is tracked over time, and we evaluate the *H_e_* distribution of 25% sliding temporal windows throughout fragmentation. As with the previous scenario, we tracked changes in the summary statistics of the distributions along fragmentation. Unlike the scenario that tracks the entire metapopulation, here we were not able to identify substantial early warning signals (Fig. 5f–h). While the standard deviation did increase as fragmentation progressed, the change was not substantial prior to the tipping-point phase (Fig. 5f). Although no directional change in kurtosis was observed, skewness showed a moderate early increase, which could potentially provide some early warning (Fig. 5g, h). Taken together, our analyses indicate that under the simulation settings examined, cross-sectional monitoring of multiple populations at each sampling occasion yields earlier and more reliable early warning signals than tracking a single population through time, even when the latter is summarized over an extended temporal window. The likely reason is that the cross-sectional snapshot includes multiple quasi-independent observations per time step, whereas the sliding-window yields serially autocorrelated records.

## Discussion

Habitat fragmentation is one of the most pressing threats to global biodiversity [2, 47], and genetic monitoring could be instrumental in tracking and managing it. However, developing monitoring and intervention strategies that take into account the real-world complexities of population structure remains a challenge [48, 49]. We present a framework that enables the modeling of habitat fragmentation and its impacts on population genetic measures, thereby expanding the potential scope of genetic monitoring. Using this framework, we model complex connectivity patterns and simulate temporal dynamics and spatially heterogeneous fragmentation processes. We examined the effects of different fragmentation scenarios on genetic measures and found that the same rate of fragmentation can lead to markedly different patterns of genetic differentiation between populations (*F_ST_*) and levels of genetic diversity within populations (*H_e_*). In this network-based perspective of fragmentation, we also find that classical population genetic relationships, such as the association between *F_ST_* and geographical distance or between gene flow and local genetic diversity, may not always hold. Network topology metrics can help interpret these associations. Finally, we demonstrate how genetic monitoring can potentially be used to detect early warning signals before fragmentation triggers critical shifts in the genetic health of populations.

The population network framework presented here can be applied to study many ecological processes that affect connectivity. Nonetheless, it is especially relevant in the context of genetic monitoring because it can inform how genetic measures in populations change over time [50]. Human activity can induce fragmentation in different ways, but theoretical investigations of fragmentation dynamics and their potential consequences have thus far been limited [21, 51]. Our results underscore the importance of considering the sequence of events leading to fragmentation for accurate evaluation of its progression. While we observe steady rates of genetic changes that are consistent with theory in some cases [52, 53], we also find scenarios in which genetic measures change abruptly (Fig. 2).

An important factor in shaping the temporal change in genetic measures is the maximum number of connected populations in the network (i.e., the size of the largest component; Fig. 3h). For example, scenarios in which a large component is maintained for a longer period (best-case, random, autocorrelated; Fig. S3) maintain the genetic health of populations longer (Fig. 2). This pattern holds even when populations within components are weakly or indirectly connected. From a landscape management perspective, it implies that enhancing connectivity between network modules (i.e., clusters of connected populations) may be more beneficial for maintaining high levels of genetic diversity than increasing direct connectivity within a weakly connected module. This result is consistent with the expectation that larger populations (or metapopulations) will exhibit higher genetic diversity due to increased gene flow and decreased genetic drift at the global scale [38]. However, increasing global connectivity can lead to homogenization of genetic pools and loss of local adaptations [54, 55]. Therefore, considering the spatial scale at which connectivity between populations is measured is crucial for accurately interpreting genetic monitoring outputs.

Populations and ecological systems facing environmental changes can undergo dramatic, unexpected, and often irreversible transitions. In the context of tracking biodiversity, several studies have introduced the concept of fragmentation thresholds that lead to regime shifts in biodiversity [56–58]. However, regime shifts in terms of genetic health and population-level metrics have received far less attention and have been considered primarily in the context of adaptive evolution in response to stress [59]. Consequently, genetic monitoring of populations is often reduced to qualitative assessments. We demonstrated that genetic indicators may appear constant during substantial periods of fragmentation, followed by rapid shifts in genetic metrics (e.g., random or autocorrelated fragmentation in Fig. 2). This suggests that a standard interpretation of genetic monitoring—no genetic change over time implies no underlying fragmentation process—can be misleading. As a proof-of-concept, we showed that early warning signals may be detectable by tracking features of the distributions of genetic monitoring data. This is particularly true if a large number of populations in the metapopulation are monitored. Although there is a substantial body of theoretical and statistical literature on early warning signals [42, 43, 46, 60], to the best of our knowledge, no theoretical or empirical studies have explored the integration of these methods with population-genetic data so far. Further investigation, applying a more comprehensive suite of early warning methods (e.g., Kendall’s *τ* statistic, conditional heteroskedasticity) to empirical data, may shed additional light on the effectiveness of this approach.

Patterns of spatial genetic structure have been extensively studied for almost a century, both in theoretical population genetic models [17, 18, 35, 61] and in empirical studies of natural populations [62–64]. One prevailing view is that spatial separation generates isolation-by-distance patterns reflected in genetic differentiation measures [17, 18, 65]. However, we find that these patterns may deviate from classical expectations depending on the underlying fragmentation scenario and the distance metric used (Fig. 4c–f). Similarly, the relationship between genetic diversity and connectivity [66–68], a key guideline in conservation practices [69], can also weaken during fragmentation (Fig. 4a–b). These findings highlight the need to integrate complex spatial configurations of populations and realistic descriptions of ecological processes into population genetic studies.

Although our framework is flexible and allows detailed spatial configurations, we assumed constant population sizes and symmetric continuous migration rates, and we did not incorporate extinction-colonization dynamics. While our sensitivity analyses suggest that the way different fragmentation scenarios affect genetic measures is relatively general and not strongly affected by the initial network structure or migration rates, other ecological features may have important impacts. Our main goal, therefore, is to provide qualitative understanding of how genetic monitoring data should be interpreted, rather than to offer precise ways to represent realistic population dynamics. One important assumption in our model relates to the time required for a system to reach migration-drift equilibrium between fragmentation steps. When the rate of fragmentation is substantially faster than the rate of approach to equilibrium, our framework may not be appropriate. It has been suggested that genetic differentiation may respond more rapidly than heterozygosity to changes in migration [52] and reach equilibrium faster [70, 71]; therefore, in some cases, the framework may be suitable for tracking genetic differentiation but not genetic diversity.

As non-invasive population-genomic data become increasingly accessible, genetic monitoring is expected to emerge as a leading tool in conservation biology for assessing the health, ecology, and behavior of natural animal and plant populations. However, the gap between theoretical expectations and practical challenges in conservation biology currently limits our ability to accurately interpret genetic data and develop landscape-specific and species-specific conservation strategies. Our framework incorporates the real-world complexities of space and time and is readily interpretable in terms of genetic monitoring. Here, we explored an important aspect of fragmentation—the processes and patterns by which between-population connectivity is lost—but our framework can be readily expanded to investigate other anthropogenic effects, such as habitat loss (e.g., by simulating different node-removal processes) or the utility of interventions (e.g., prioritization of ecological corridors). Our network-based framework thus serves to narrow the gap between theoretical insights and the complex ecological realities of conservation biology.

## Methods

All analyses were performed using Python 3.11.1, except where stated otherwise.

### Computing genetic measures in population networks

To compute genetic measures for population networks, we employed the framework developed by Alcala *et al.* [34], which integrates the mathematical relationship between migration and coalescence times by Wilkinson-Herbots [35] with the relationship between coalescent times and *F_ST_* by Slatkin [36]. Our method relies on transformations among three matrices: (i) the migration matrix describing the pairwise migration rates, (ii) the coalescence matrix describing the expected time to coalesce for two lineages within or between populations, and (iii) the *F_ST_* matrix describing the pairwise genetic distance between populations. A full explanation of the derivations and computations is presented in the Supplementary Information Text.

We considered an idealized system of *K* populations of equal size, evolving under the neutral Wright-Fisher model at migration-drift equilibrium [19]. Let *m_ij_* denote the backward migration rate from population *i* to *j*, representing the probability that a lineage in *i* originated in *j* in the previous generation. We assumed symmetric migration (*m_ij_* = *m_ji_* for all *i* and *j*) to ensure conservative migration [35], where total incoming and outgoing migration balance in each population: Σ*_j_*_≠*i*_ *M_ij_* = Σ*_j_*_≠*i*_ *M_ji_*. While conservative migration is a weaker assumption than symmetric migration, we imposed symmetric migration for tractability. Under these assumptions, the migration structure of the populations is represented as a symmetric, undirected network *M* of *K* nodes with zero-diagonal entries. For a pair of nodes *i* and *j* (*i* ≠ *j*), the weight assigned to the edge is *M_ij_* = 4*Nm_ij_*, representing the expected number of migrants from *i* to *j* per generation, with *N* denoting the population size of each of the nodes. We simulated population networks with *K* = 50 nodes, where migration rates are uniform across all edges (*M_ij_* = 1 in the main text, and alternative migration rates in Fig. S1).

### Simulating fragmentation processes in population networks

Because natural populations are embedded in a geographic space, we used spatial network models [72], in which nodes correspond to populations with assigned geographic coordinates. We primarily used the random geometric graph (RGG) model [37], one of the simplest and most widely studied spatial network models, to generate the initial network in our simulations (see Fig. S2 for alternative network models). In this model, *K* populations are placed uniformly at random in a unit square in Euclidean space, and an edge is formed between two nodes if their Euclidean distance is below a fixed threshold *d*. The RGG model is particularly well-suited for representing migration in spatially structured populations because it captures the ecologically realistic constraint that migration occurs only between sufficiently proximate populations. The connectivity threshold for two-dimensional RGG networks (i.e., the value of *d* above which the network is almost surely connected) is 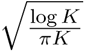 [73], which equals *d* = 0.16 for *K* = 50. We therefore set *d* = 0.30, which consistently generates a connected network that is not too dense yet sufficiently above the threshold at which the network is close to being disconnected.

To model the fragmentation process, we sequentially remove edges from the initial network, one at a time, until no edges remain. We consider eight fragmentation scenarios (Fig. 1). (i) *Random fragmentation*. At each fragmentation step, an edge is removed uniformly at random, representing non-specific habitat deterioration, such as fragmentation induced by global climate change. (ii) *Autocorrelated fragmentation*. Initially, one random edge is removed. At each subsequent step, one edge is removed uniformly at random from the set of edges adjacent to the previously removed edge (i.e., edges sharing a node with the last removed edge). This process models spatially correlated landscape disturbances, such as urban or agricultural expansions. (iii) *Intrusive fragmentation*. A node is selected uniformly at random, and all its incident edges are removed in random order. Once these edges are removed, another node is chosen randomly, its incident edges are removed, and the process is repeated. This process generates isolated habitable “islands” within the landscape, representing, for example, the formation of micro-reserves—small, disconnected populations. (iv) *Regressive fragmentation*. Edges are sorted by the minimum x-coordinate of their incident nodes in the Euclidean plane and removed progressively from low to high x-coordinate values, starting with the edge having the smallest x-coordinate. This process represents large-scale spatial disturbances moving across the habitat, such as shifts in climate-change fronts. (v) *Distance-based fragmentation*. At each step, the edge connecting the most distant populations in the underlying Euclidean space is removed. This process represents a general environmental deterioration that impedes long-distance dispersal among habitat patches. (vi) *Divisive fragmentation*. A line is drawn in the Euclidean plane by connecting two points on different boundaries (either opposing or neighboring boundaries) of the metric space (selected uniformly at random), effectively bisecting the habitat. All edges intersecting this line are sequentially removed, starting with those having the smallest x-coordinate (as defined in (iv)). This process models the introduction of linear barriers, such as roads or railways, into the landscape. (vii) *Best-case fragmentation*. At each step, the edge with the lowest betweenness centrality is removed. Betweenness centrality was computed with the NetworkX Python library. Because such edges contribute minimally to network connectivity, removing them is expected to have the least impact on genetic measures. Although this scenario is not realistic, it serves as an upper benchmark for evaluating genetic measures at a given level of fragmentation. (viii) *Worst-case fragmentation*. Similar to best-case fragmentation, but at each step, the edge with the highest betweenness centrality is removed. This process provides a lower benchmark for genetic measures at a given level of fragmentation.

These eight fragmentation processes do not exhaustively cover all possible scenarios, but rather describe typical ecological and anthropogenic disturbance patterns relevant to genetic monitoring [21, 74]. Because these processes are stochastic, we performed 100 independent replicates per fragmentation type, randomizing the initial network configuration and the fragmentation sequence in each replicate.

In each simulation replicate, we computed the changes in *F_ST_* and *H_e_* distributions in response to fragmentation, assuming migration-drift equilibrium is reached between successive iterations of edge removal. Each replicate generates a sequence of migration matrices *M*_0_*, …, M_x_*, with *x* being the last fragmentation step. From these migration matrices, we computed corresponding *F_ST_* matrices *F*_0_*, …, F_x_* and *H_e_* vectors *H*_0_*, …, H_x_*. These sequences reflect the changes in genetic differentiation and genetic diversity throughout fragmentation. Using these sequences, we tracked changes in the means (Fig. 2), sample variances (Fig. 3g) and distributions (Fig. 3a–f) of the genetic measures along fragmentation.

We also evaluated changes in network structure throughout fragmentation by tracking for four structural categories: (i) largest component, (ii) other components with *>* 3 populations (medium components), (iii) components of 2–3 populations (pairs/triads), and (iv) isolated nodes. At each time step, we computed the mean proportion of nodes in each category across simulation replicates.

To account for alternative patterns of gene flow in our initial network, and to evaluate their effect on our main conclusions, we considered two additional models. (i) The Erdős–Ŕenyi (ER) random network [75], in which, for *K* populations, each pair of populations is connected by an edge with probability *p*. To generate a well-but not fully-connected initial network, we set *p* = 0.1 (125 edges in total). To allow spatially explicit analysis that can be compared to the RGG, we embedded this model in Euclidean space, with nodes placed uniformly at random. (ii) The small-world Watts–Strogatz (WS) network is constructed by connecting each population (node) to its *k* nearest neighbors in a ring topology, and then rewiring each edge with probability *p* to connect to a randomly chosen population (node), introducing long-range connections while preserving the total number of edges. The WS network can represent species with a life history of many short-distance dispersal events and an occasional long-distance dispersal event. We use a modified variant of this model to incorporate spatial characteristics to the network [76, 77]. We use the grid graph function in the Python library NetworkX to generate a two-dimensional network with *n* nodes, setting *k* = 4 (4 neighbors for each node), and a re-wiring probability of *p*. This setting converges to the stepping stone model [17] for *p* = 0. For our simulations, we set *n* = 49 (a 7 × 7 matrix).

### Correlations between genetic measures and node attributes

To investigate how network metrics influence genetic monitoring along fragmentation, we examined the relationship between genetic measures and network metrics. For each network at each fragmentation step, we computed two node centrality measures, degree centrality (number of incident edges for the focal node) and betweenness centrality (how often a node lies on the shortest paths between other pairs of nodes), for all nodes in the network. We then computed the Pearson correlation coefficient (*r*) between node’s *H_e_* and their centrality score, at each fragmentation step and for each centrality measure (we excluded isolated nodes, for which centrality is undefined). Then, we computed the mean *r* and its *SD* for each fragmentation step across the simulation replicates, for each one of the centrality metrics and each fragmentation scenario. We only show significant correlation results (*p <* 0.05) with data from 5 or more replicates.

Similarly, we evaluated the relationships between network distance metrics and pairwise *F_ST_*. We computed the distance between all pairs of nodes in each fragmentation step using three distance metrics: (i) Euclidean distance, the standard geometric measure in the embedded metric space, analogous to the typical geographic distances among populations; (ii) shortest-path distance [78], calculated as the minimum number of edges needed to traverse from one node to another, reflecting topology-aware movement; (iii) random-walk distance [79], defined as the mean number of edges a random walker requires to travel from one node to another, which is suitable for movement that is unaware of the network topology or a non-targeted movement [22]. Random-walk distance was estimated using 50 random-walk iterations per pairwise comparison. The correlations were calculated only within connected components of size *>* 3, and pairs of nodes in disconnected components were excluded from correlation calculations (these pairs have *F_ST_* = 1 and are infinitely distant from each other for shortest-path and random walk distance). We computed the Pearson correlation coefficient (*r*) between the *F_ST_* of all node pairs and their distance score, at each fragmentation step and for each distance metric. We used the mantel python library to perform a Mantel test and calculate a corresponding p-value with 999 permutations. For networks with multiple components, and hence multiple *r* and *p*-values, we calculated the weighted mean *r* and *p* based on the component size. We then computed the mean *r* and its SD for each fragmentation step across the simulation replicates, for each one of the distance metrics and each fragmentation scenario. We only show significant correlation results (*p <* 0.05) with data from 5 or more replicates.

### Detecting early warning signals before population collapse

To identify early warning signals, we computed several summary statistics of the genetic diversity (*H_e_*) distributions that are commonly used as early warning signals: standard deviation, skewness, and kurtosis. As the process approaches the tipping-point phase, the theoretical expectation is that the standard deviation of the *H_e_* distribution will increase, the skewness will shift toward lower *H_e_* values (higher asymmetry), and increased frequency of extreme values will lead to higher kurtosis [43, 45]. We did not use the lag-1 autocorrelation, although it is often used metric to measure the return rate to equilibrium after a perturbation [44], because this statistic it is designed to measure stability around a single equilibrium [44, 46], while our framework considers a series of fragmentation events between each the system arrives at migration-drift equilibrium.

For this analysis, we focused on autocorrelated fragmentation (Fig. 2b). We used a more connected initial network than used in previous analyses, an RGG with *d* = 0.6, to capture a substantial period that is far from the tipping-point phase. We ran 1000 simulations replicates and we considered two monitoring scenarios: (i) *entire metapopulation monitoring*, where we analyze the *H_e_* distribution across all populations in the network at each step, and (ii) *single population monitoring*, focusing on the *H_e_* of the final nodes to become isolated. In the latter case, we used the generic_ews function from the R package earlywarnings to apply a sliding window approach over time, with window size of 25% and default parameters without detrending or preprocessing the data.

## Data Availability

All code is available in the GitHub repository at https://github.com/Greenbaum-Lab/fragmentation.git

## Acknowledgments

We thank Keith Harris and Eyal Halutz for their input and assistance in the analysis. The research was funded by the Binational Science Foundation (BSF) Grant No. 2021244. OP was supported by fellowships from the Israel Ministry of Environmental Protection and from the Minerva Center for the study of Population Fragmentation (MCPF).

