## Supplementary Information for "Network-based genetic monitoring of landscape fragmentation"

### Table of contents

**Deriving genetic measures for population networks** . . . . . . . . . . . . . . . . . . . . **2**

**Supplementary figures** . . . . . . . . . . . . . . . . . . . . . . . . . . . . . . . . . . . . **4**

### SI Methods: Deriving genetic measures for population networks

We begin by formulating a transformation from a migration network $M$ with $K$ nodes to a coalescent-time matrix $T$, which is a symmetric matrix with positive entries. Entry $T_{ij}$ denotes the expected coalescence time of a pair of individuals sampled from populations $i$ and $j$, and $T_{ii}$ denotes the expected coalescence time of a pair of individuals sampled from the same population. We assume that all populations are of the same size $N$, and we scale time accordingly to coalescent units of $2N$ generations. Under these notations, the transformation from $M$ to $T$ can be computed by solving a set of linear equations [1]:

$$\begin{cases} (1+M_{ii})T_{ii} - \sum_{k\neq i} M_{ik}T_{ik} = 1, \\ \frac{1}{2}(M_{ii}+M_{jj})T_{ij} - \sum_{k\neq i} \frac{M_{ik}}{2}T_{jk} - \sum_{k\neq i} \frac{M_{jk}}{2}T_{ik} = 1, \end{cases} \tag{1}$$

with $i \neq j$ and $M_{ii} = -\sum_{k\neq i} M_{ik}$ for $i = 1, ..., K$. Eq. 1 is defined if all $T_{ij}$ are finite, which is true if and only if the migration network is connected (i.e., consists of a single component). When $M$ can be decomposed into several network components, Eq. 1 is defined within each component.

Using Eq. 1, we track the genetic diversity of each population and compute its expected heterozygosity [2, 3]:

$$H_i = (\theta_i T_{ii})/K, \tag{2}$$

where $\theta_i = 4N\mu$ is the scaled mutation rate per site per generation under the infinite-site model [3]. Because we assume equal population size and a fixed mutation rate for all populations, we simplify the notation and study the 'unscaled expected heterozygosity' $H_i = T_{ii}/K$. In other words, heterozygosity is the coalescence time for two individuals within each population normalized by the total number of populations in the network. In our model, $H_i$ of individual populations can exceed values of 1, usually in populations that act as central gene flow bridges (i.e., hubs) in the network. This is because the probability of two individuals in the central population to coalesce decreases, while the probability of the individuals to drift apart from one another increases. This pattern contrasts with the fully connected island model, where individuals are always within one migration event for each pair of populations. Thus, the interpretation of heterozygosity in our context should be interpreted only in relative values.

Next, we formulate a transformation from the coalescent times matrix $T$ to the $F_{ST}$ matrix $F$, whose entry $F_{ij}$ is a pairwise $F_{ST}$ [4] value between populations $i$ and $j$. For low mutation rates, $F$ can be approximated from $T$ using a set of non-linear equations [2, 5]:

$$F_{ij} = \frac{T_T^{ij} - T_S^{ij}}{T_T^{ij}}, \tag{3}$$

where $T_S^{ij} = (T_{ii} + T_{jj})/2$ is the expected within-population coalescence time, and $T_T^{ij} = (T_{ij} +$

$T_S^{ij})/2$ is the expected coalescence time of two individuals sampled from these two populations. By definition, $F$ is symmetric, with zero diagonal entries. Eq. 3 is derived under the assumption of small mutation rates, which was shown to be valid under many realistic scenarios [6]. Following Eq. 1, Eq. 3 is also defined only for populations in the same component of $M$. For two populations $i$ and $j$ in different components of $M$, we set $F_{ij} = 1$ because there is no gene flow between the populations and they are therefore maximally differentiated.

### Supplementary Figures

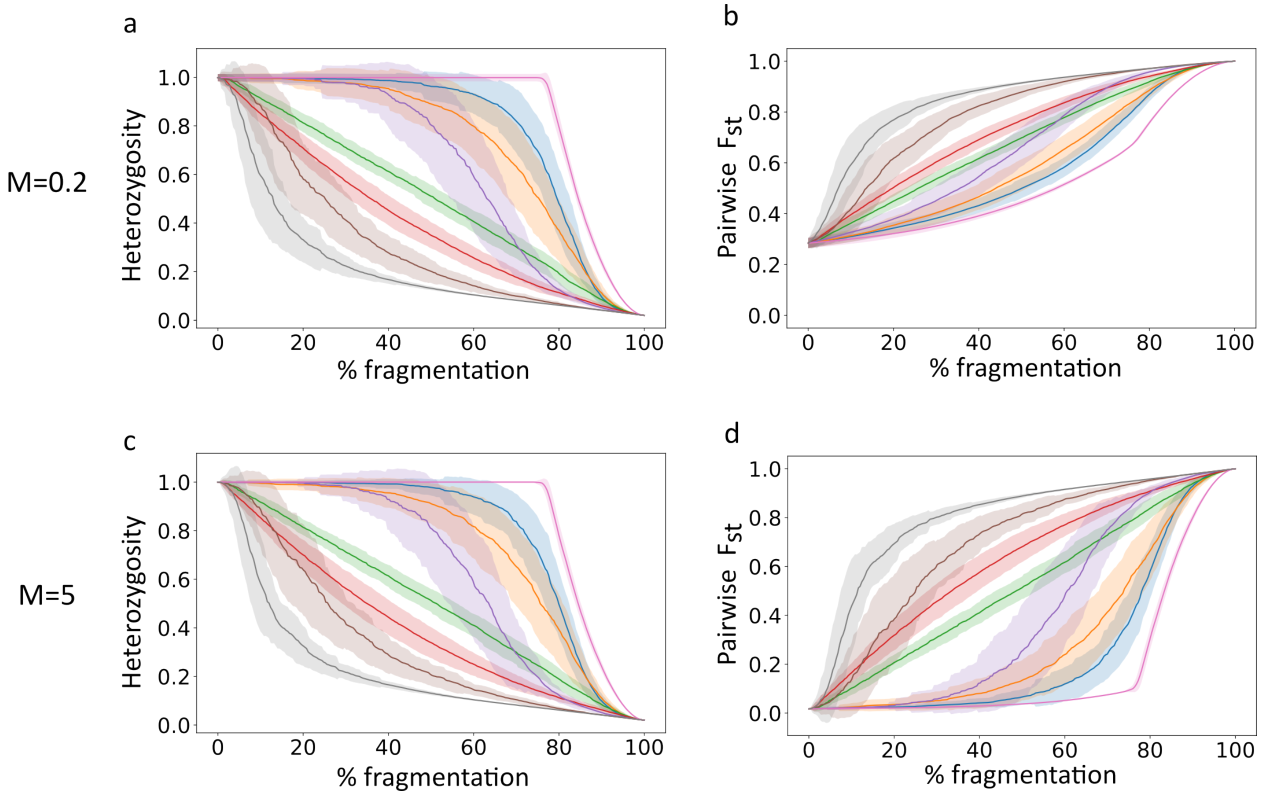


Figure S1: **Changes in genetic measures during fragmentation under alternative migration rates (M).** (a–b) Mean heterozygosity and $F_{ST}$ under low migration rates $M = 0.2$. (c–d) Mean heterozygosity and $F_{ST}$ under high migration rates $M = 5$. The curves denote the mean across 100 simulation replicates, and the shaded areas denote the standard deviation. Colors as in Fig. 2.

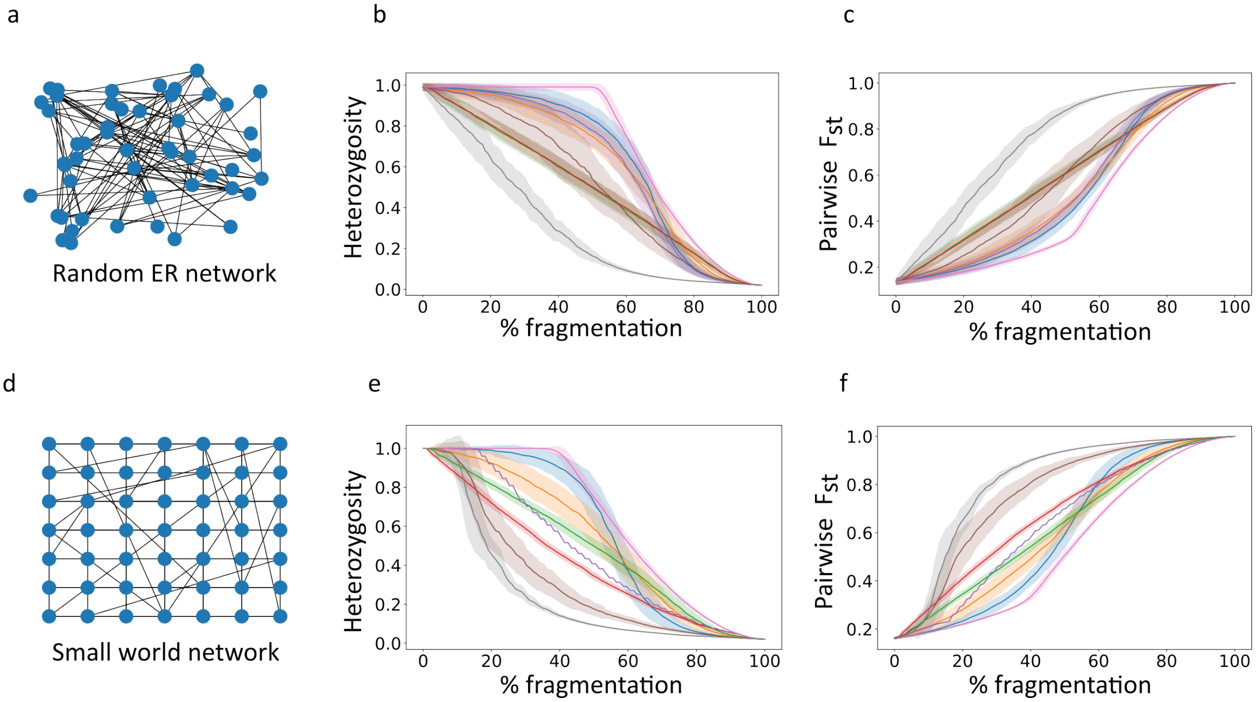


Figure S2: **Changes in genetic measures along fragmentation for alternative initial networks.** (a) Example of an ER network realization in which nodes are connected to each other randomly at uniform with probability $p = 0.3$. (b–c) Mean heterozygosity and $F_{ST}$ for ER network. (d) Example of a spatial small-world network realization in which nodes are connected to their closest neighbors with additional random connections at probability $p = 0.015$. (e–f) Mean heterozygosity and $F_{ST}$ for small-world network. The curves denote the mean across 100 simulation replicates, and the shaded areas denote the standard deviation. Colors as in Fig. 2.

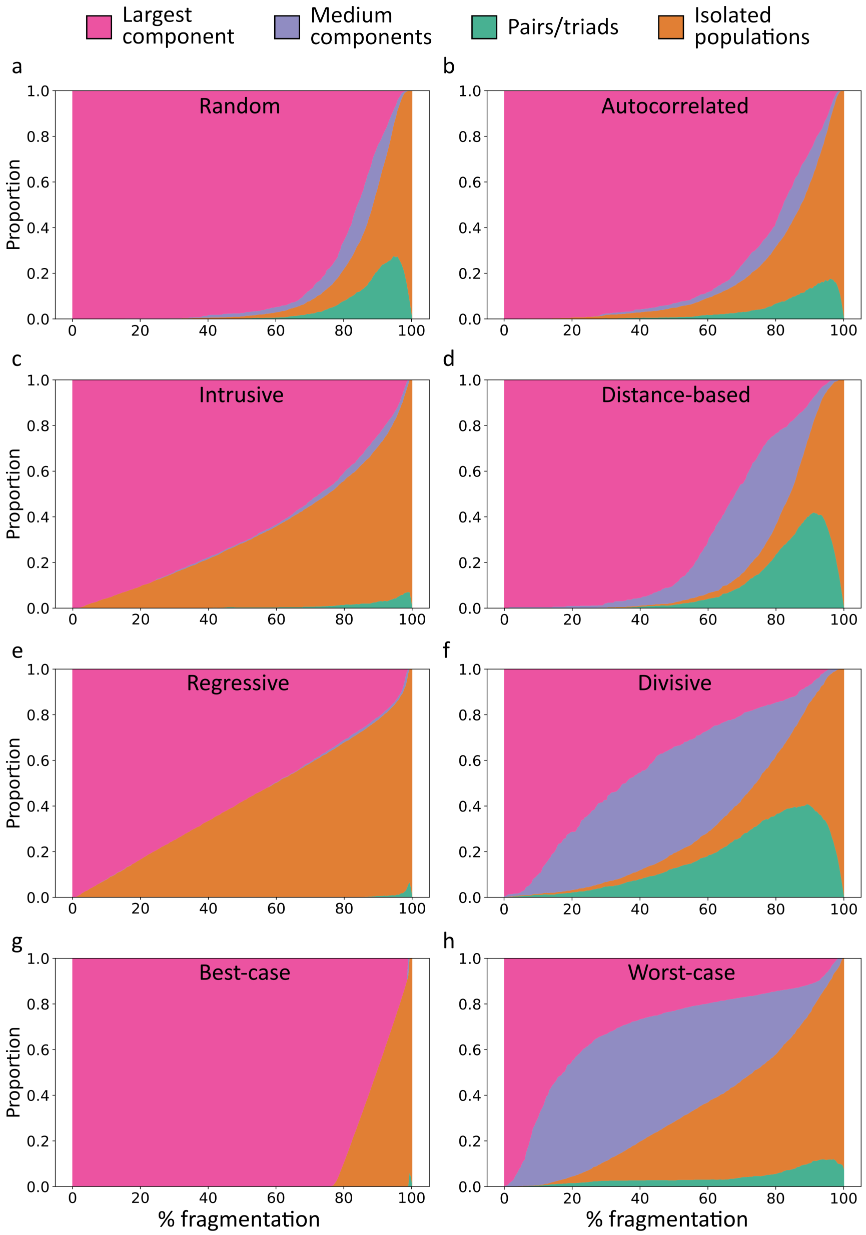


Figure S3: **The proportions of different network structures throughout eight fragmentation scenarios.** Largest component–the largest set of connected nodes in the network; medium components–a connected set of >= 4 nodes, excluding the largest component; pairs/triads–two or three connected nodes; isolated populations–nodes without edges. For each scenario we show the mean across all simulation replicates.

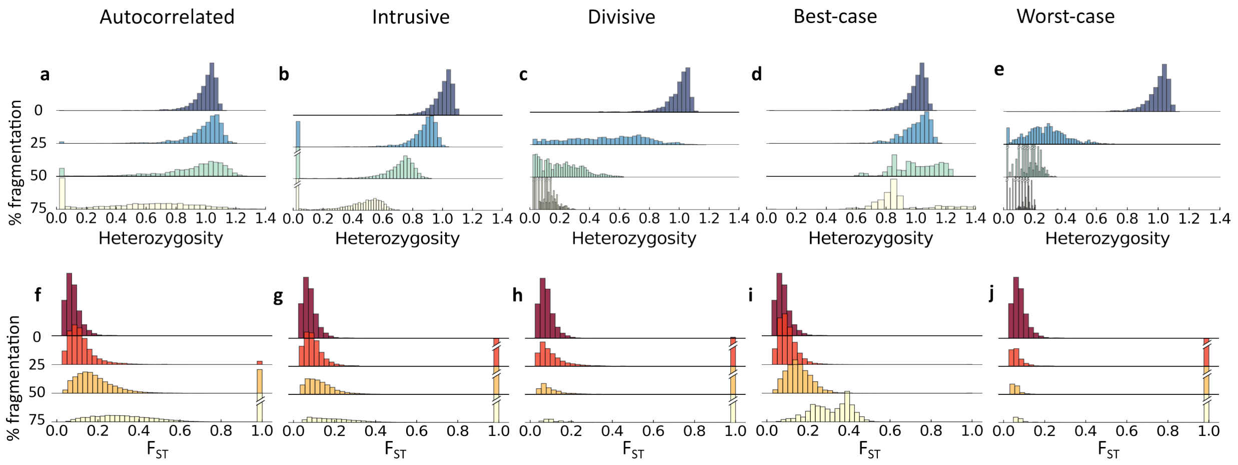


Figure S4: **Changes in distribution of genetic measures along fragmentation.** Presented here are the remaining five fragmentation scenarios that are not included in the main text: autocorrelated, intrusive, divisive, best-case, and worst-case. Shown are four snapshots during the process, with 0, 25, 50, and 75% of the edges removed. Diagonal lines on bars represent high values that were truncated for better visualization. (a–e) Distribution of expected heterozygosity ($H_e$) within populations. (f–j) Distribution of pairwise $F_{ST}$ values. $F_{ST} = 1$ indicates pairs of nodes that are not in different network component (i.e., not connected by any path). In all panels, distributions are pooled across 100 simulation replicates.

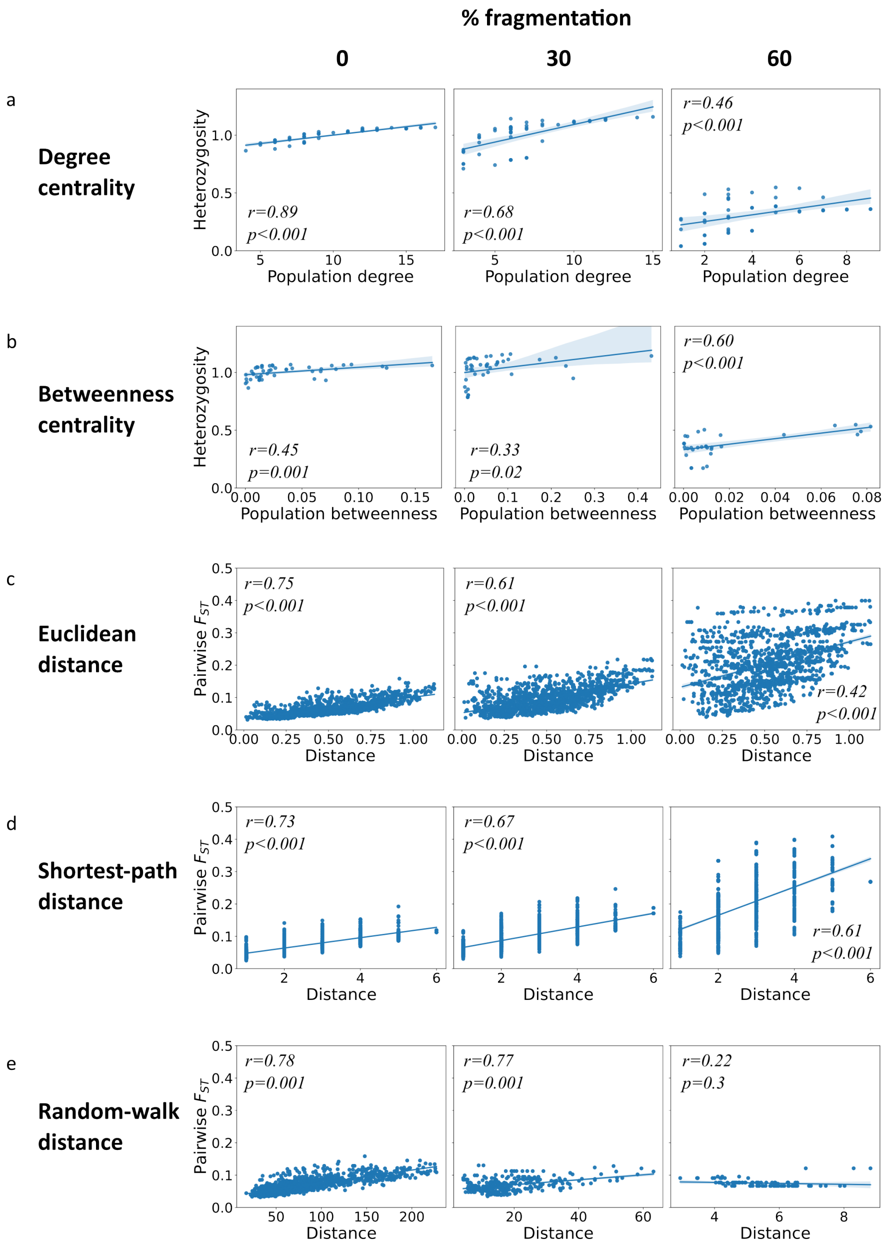


Figure S5: **Snapshots of correlation between population genetic measures and network metrics.** The Regression plots show snapshots at 0%, 30%, and 60% level of fragmentation. $r$ and $p - value$ are shown inline. (a–b) Correlation under distance-based scenario, (c–d) Correlation under best-case scenario. (e) Correlation under worst-case scenario.
